## Supplementary Figures and Tables for "A simple method for developing lysine targeted covalent protein reagents"

### Supplementary information

#### Supplementary Figures

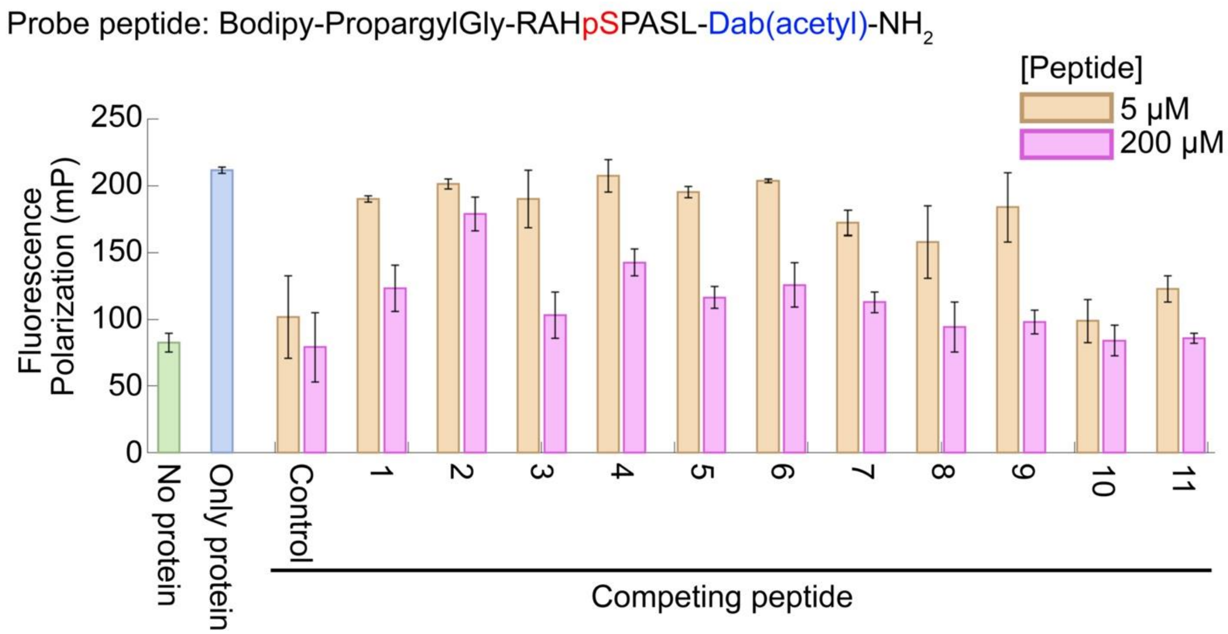

**Supplementary Figure 1. Fluorescence Polarization competition experiments.**
Fluorescence polarization of 5 nM BODIPY-labeled noncovalent binder for 14-3-3σ was measured alone (green), in the presence of 0.25 μM 14-3-3σ (blue) and in the presence of 14-3-3σ and the different electrophilic peptides (orange and magenta). The sequence of the BDP labeled binder is shown on top, Dab = diaminobutyric acid. The control peptide is identical to the BDP-labeled binder in sequence.

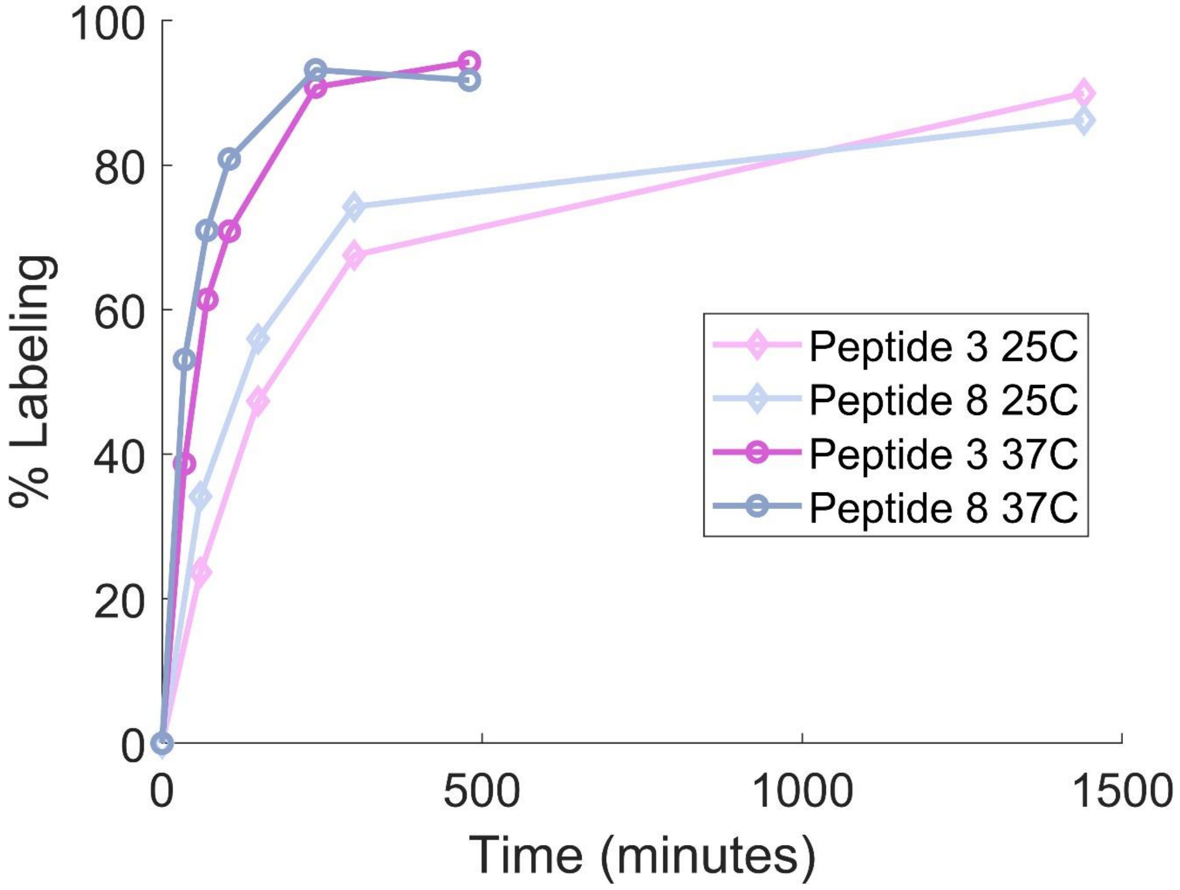

**Supplementary Figure 2. Peptide labeling at various temperatures.**
Peptides **3** and **8** (5 µM) were incubated with 14-3-3σ (2 µM) at either 25°C or 37°C and the degree of labeling was measured using LC/MS.

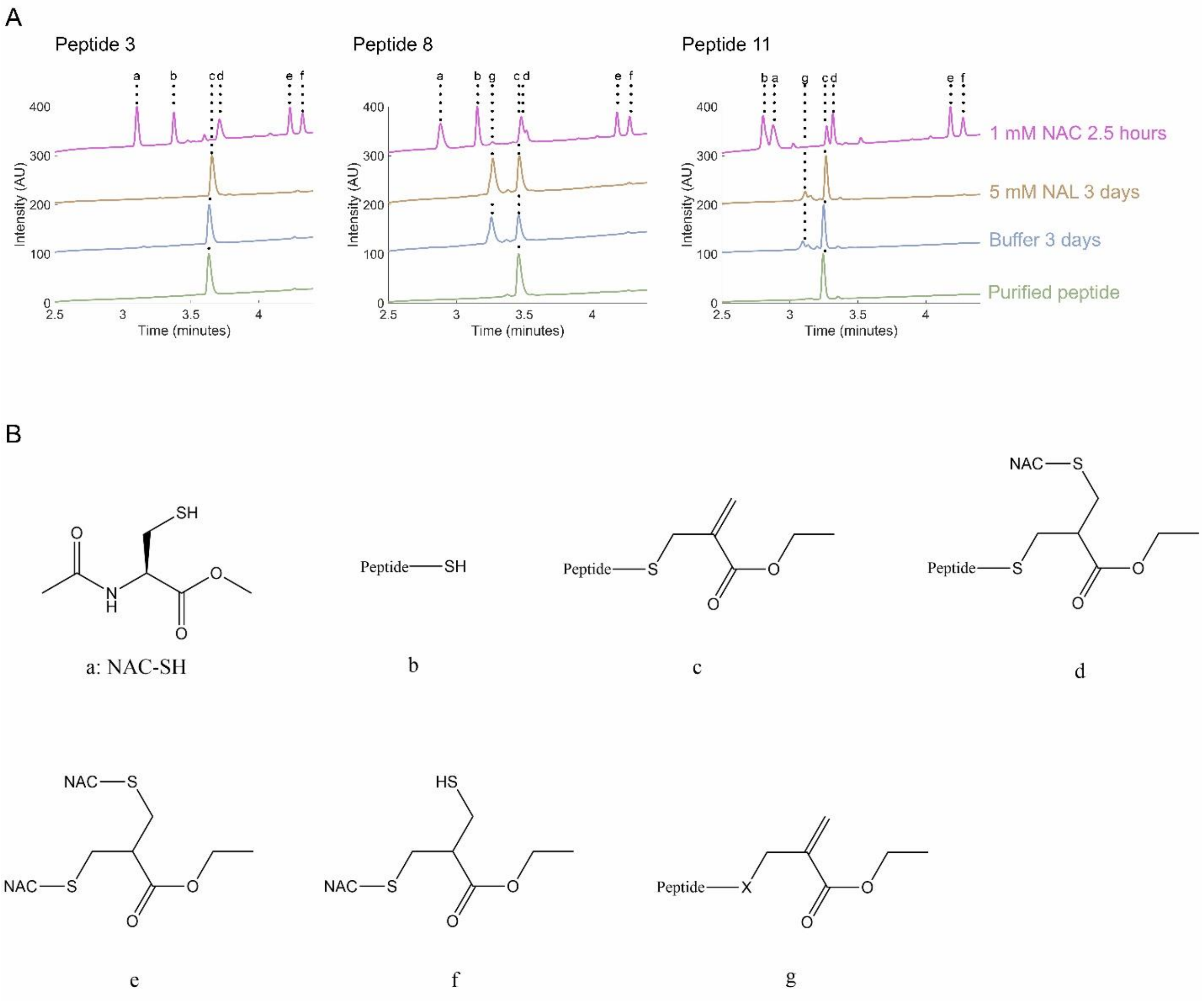

**Supplementary Figure 3. Peptide stability in buffer and in the presence of lysine and cysteine.**
**A.** Peptides **3**, **8** and **11** were incubated with 1 mM N-acetyl cysteine methyl ester (NAC, magenta) for 2.5 hours, 5 mM N-acetyl-lysine methylester (NAL, gold) for 3 days, and buffer only (HEPES 25 mM pH = 7.5, 100 mM NaCl, 10 mM $MgCl_2$) for 3 days, and the products were analyzed using LC/MS. The products were identified as shown in panel **B.** Product **g** is an isomerization product of peptides **8** and **11**, possibly due to migration of the methacrylate to a different position in the peptide, as it shows the same molecular mass, but different retention times to product **c**.

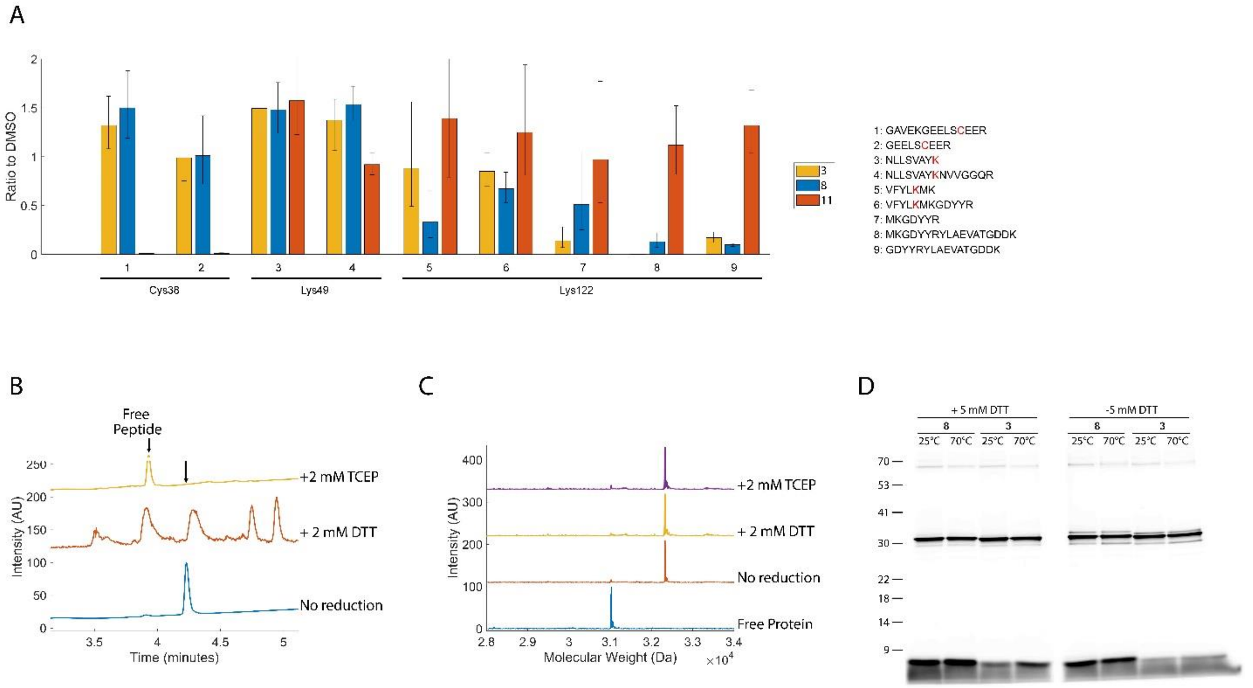

**Supplementary Figure 4. Methacrylate peptides label different 14-3-3σ residues.**

**A.** Following incubation with the peptides, samples of 14-3-3σ were trypsinized and tryptic peptides containing or following potential target residues were quantified using parallel reaction monitoring. **B.** Peptide **3** was incubated with 2 mM of either TCEP or DTT for 130 minutes at room temperature and the products were characterized using LC/MS. **C.** 14-3-3σ was incubated with peptide **3** until fully labeled, followed by incubation with TCEP or DTT for 130 minutes at room temperature and analysis by LC/MS. **D.** 14-3-3σ was incubated with BODIPY-modified peptides until fully labeled. The samples were then denatured using Lithium Dodecyl Sulfate buffer in different conditions (with or without heating and with or without DTT) and analyzed by SDS-PAGE.

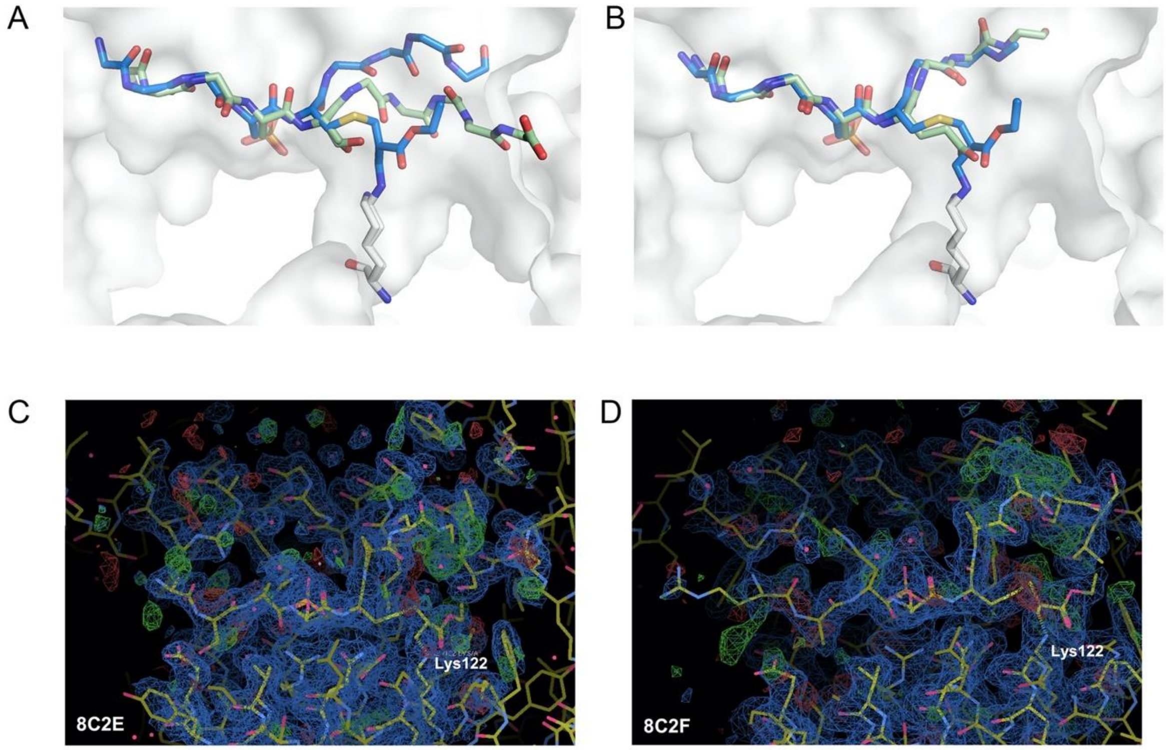

**Supplementary Figure 5. Comparison of structures of covalent complexes and noncovalent complexes.**
**A.** Comparison of the structure of the covalent complex between peptide **8** and 14-3-3σ (blue) with the noncovalent complex with the parent peptide (green). **B.** Comparison of the structure of the covalent complex between peptide **3** and 14-3-3σ (blue) with the noncovalent complex with the parent peptide (green). **C.** Portion of the electron density maps for peptide **3.** Displayed are 2Fo-Fc (blue mesh, contoured at 1.0 σ) and Fo-Fc maps (green and red meshes, contoured at 2.5 σ). **D.** Portion of the electron density maps for peptide **8.** Displayed are 2Fo-Fc (blue mesh, contoured at 1.0 σ) and Fo-Fc maps (green and red meshes, contoured at 2.5 σ).

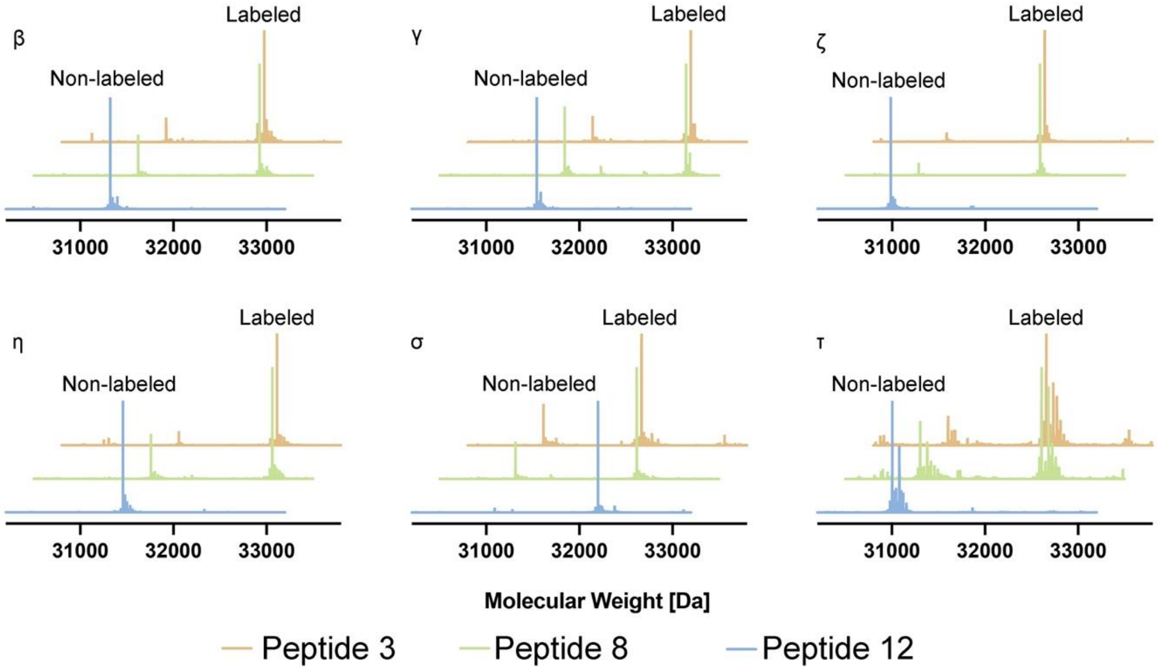

**Supplementary Figure 6: Isoform selectivity of 14-3-3 covalent binding peptides**

Binding of electrophilic peptides to 14-3-3 isoforms. Peptides were incubated with the various indicated isoforms overnight at room temperature, diluted and analyzed using intact protein LC/MS.

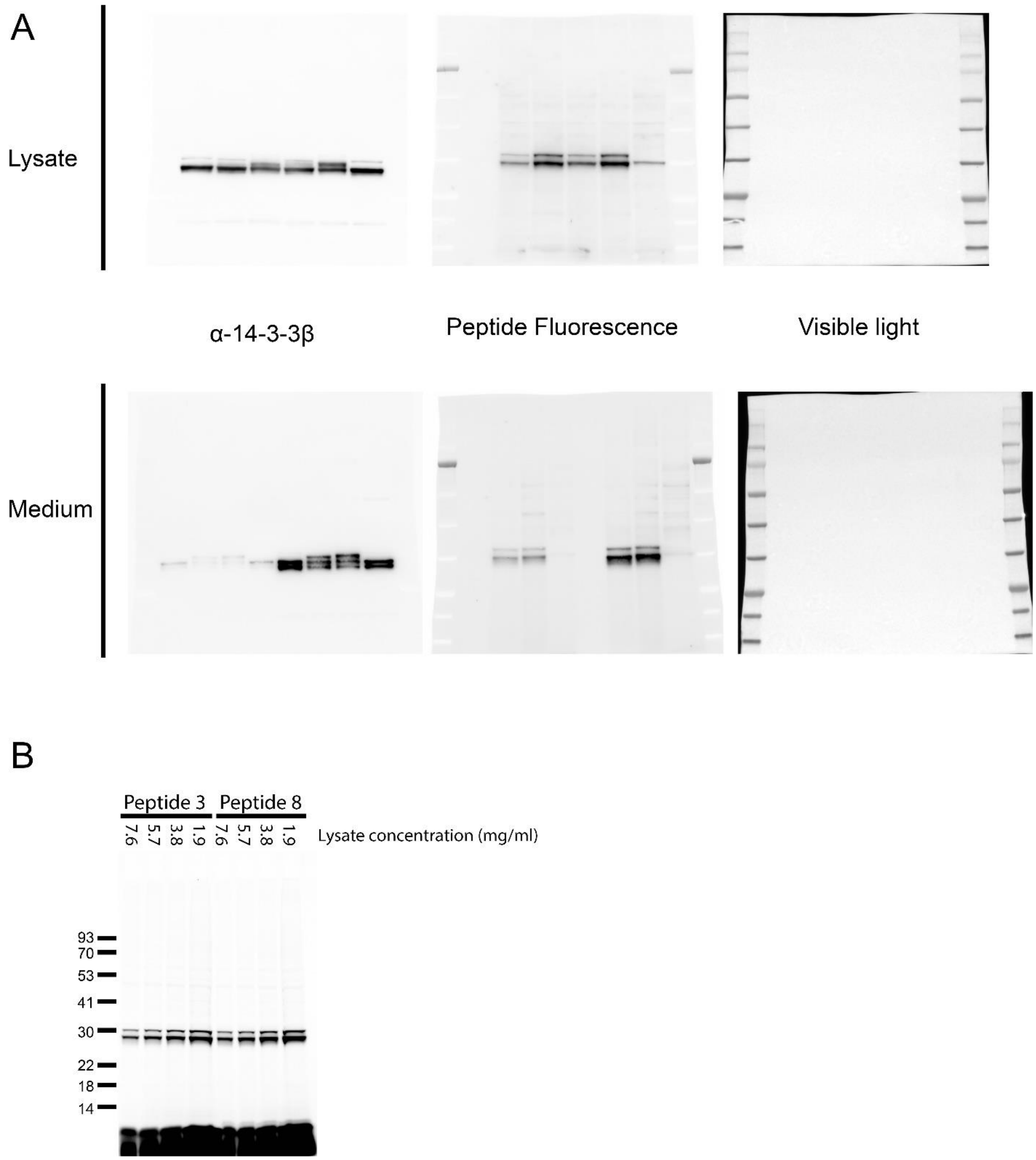

**Supplementary Figure 7. Peptide binding selectivity in cell lysates and media.**
**A.** Raw images for the western blot experiment depicted in figure 5. Experiments was repeated 3 times. Top row: images for experiment done on A549 lysates. Bottom row: Images from extracellular medium. **B.** Direct gel imaging for experiment in which 1 µM peptide was incubated with A549 lysates at the indicated concentrations for 22 hours at 25°C. The same amount of lysate was loaded in each lane (28.5 µg). The intense fluorescence at the bottom comes from unbound peptide.

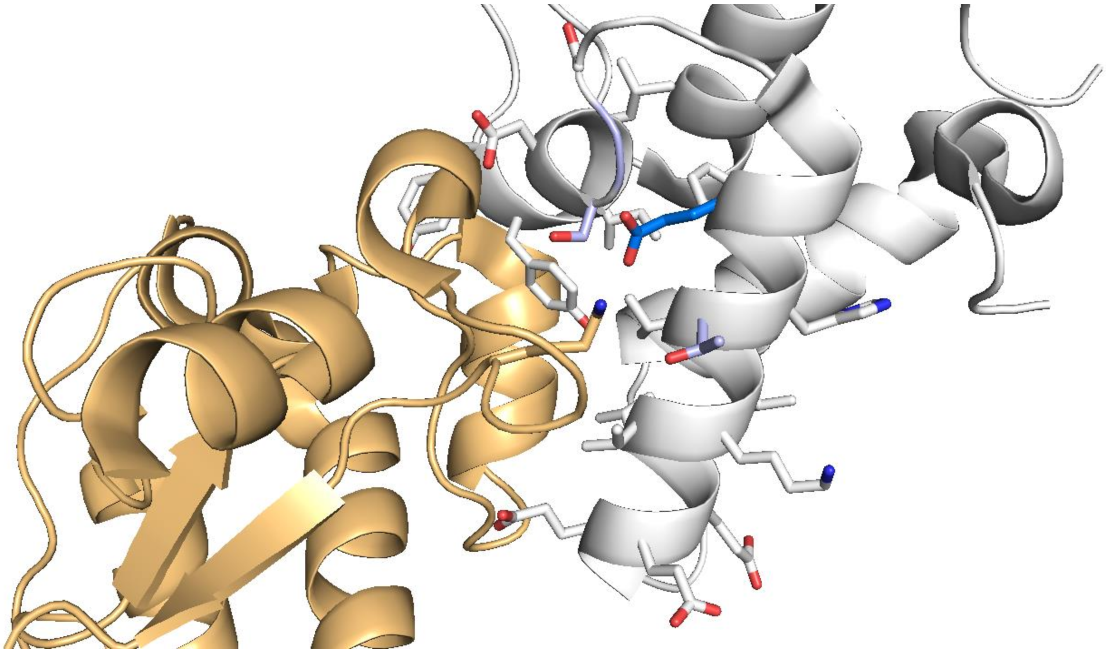

**Supplementary Figure 8. Structure of the Im9-E9 complex (Im9: white; E9: gold).**
Im9 residues mutated to cysteine and modeled as methacrylates are shown as white sticks. Residues that were mutated and gave interface RMSD below 1 Å are depicted in light blue. The selected residue (Glu41) is shown in blue, while the target residue in E9 (Lys122) is shown in gold.

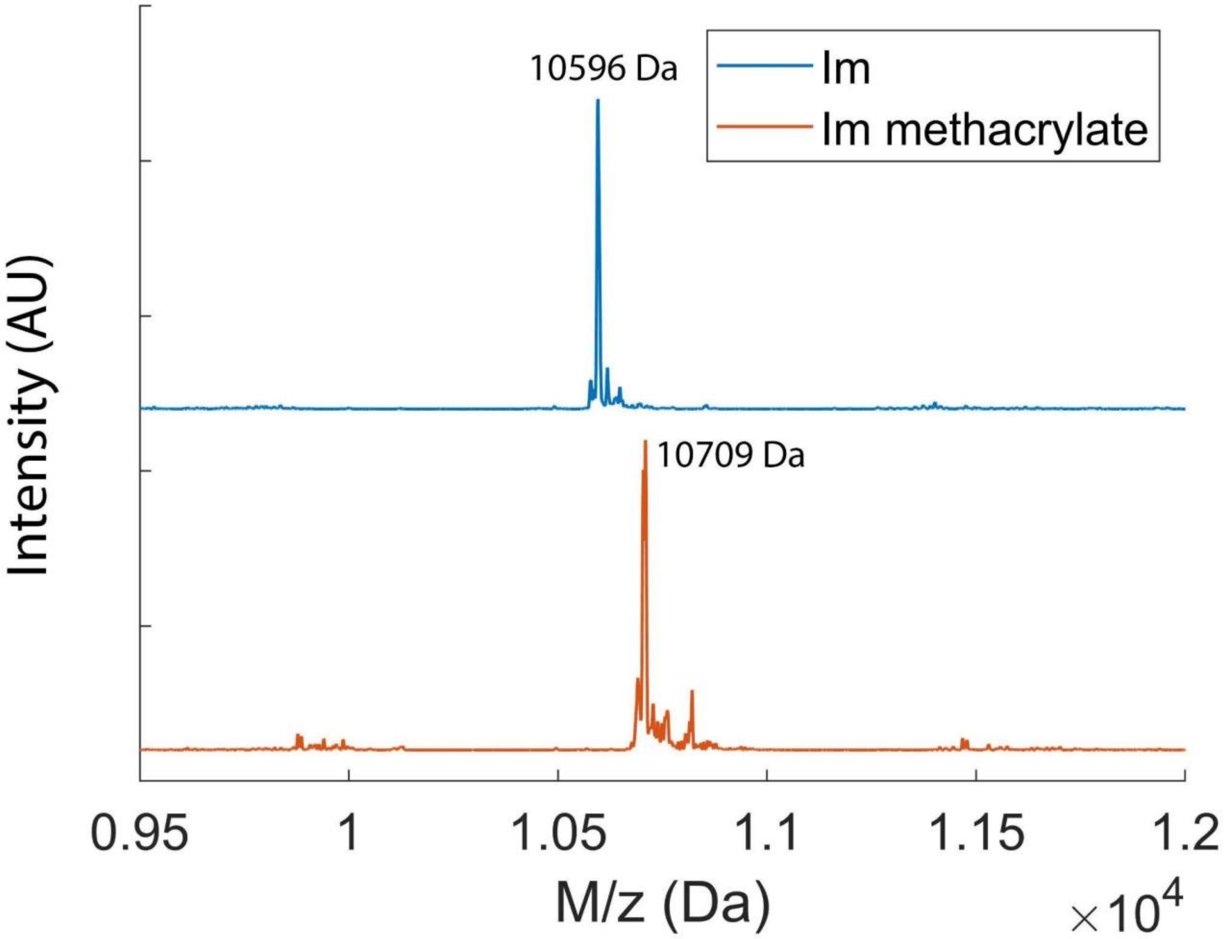

**Supplementary Figure 9.** Deconvoluted mass spectra of Im C23A/E41C mutant (Top) and the methacrylate-modified protein (bottom).

A

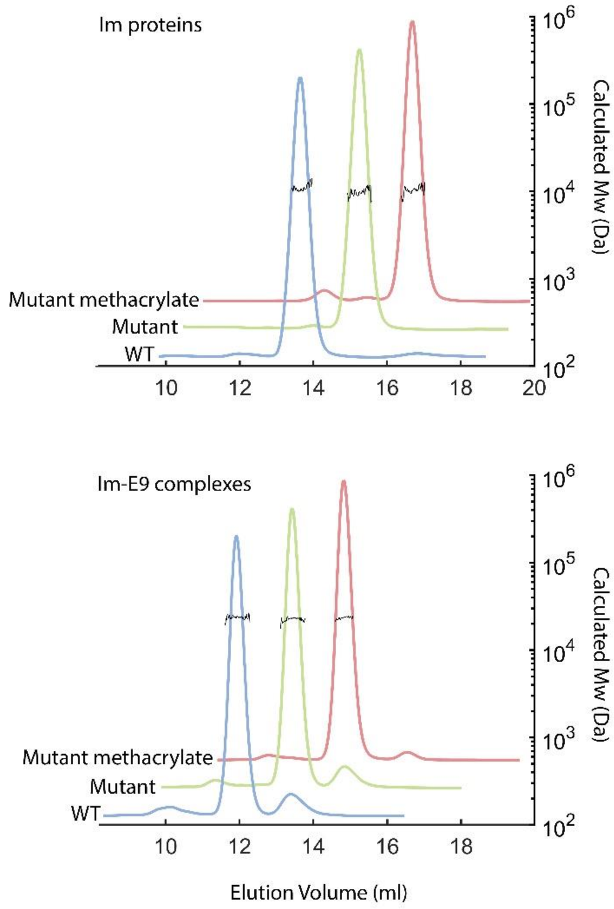

B

| Protein | Mw from elution volume (kDa) | Mw from MALS (kDa) |
|---|---|---|
| **Im** | **11.5 ± 0.8** | **11.0 ± 0.2** |
| **Im-mut** | **9.7 ± 1.3** | **9.92 ± 0.17** |
| **Im-mut methacrylate** | **10.3 ± 0.7** | **10.6 ± 0.2** |
| **Im-E9 complex** | **24.0 ± 1.0** | **23.7 ± 0.2** |
| **Im-mut-E9 complex** | **22.6 ± 0.7** | **22.8 ± 0.2** |
| **Covalent complex** | **23.4 ± 0.7** | **23.6 ± 0.1** |

**Supplementary Figure 10. SEC-MALS evaluation of Im9/E9 complexes.**

**A.** SEC-MALS measurements of Im proteins alone (top) and in complex with E9 (bottom), measured in Phosphate Buffered Saline, pH = 7.4. The colored plots show the normalized UV absorbance, and the black plots indicate the measured molecular weight from MALS. **B.** Calculated molecular weights of the constructs from both elution volume and MALS data.

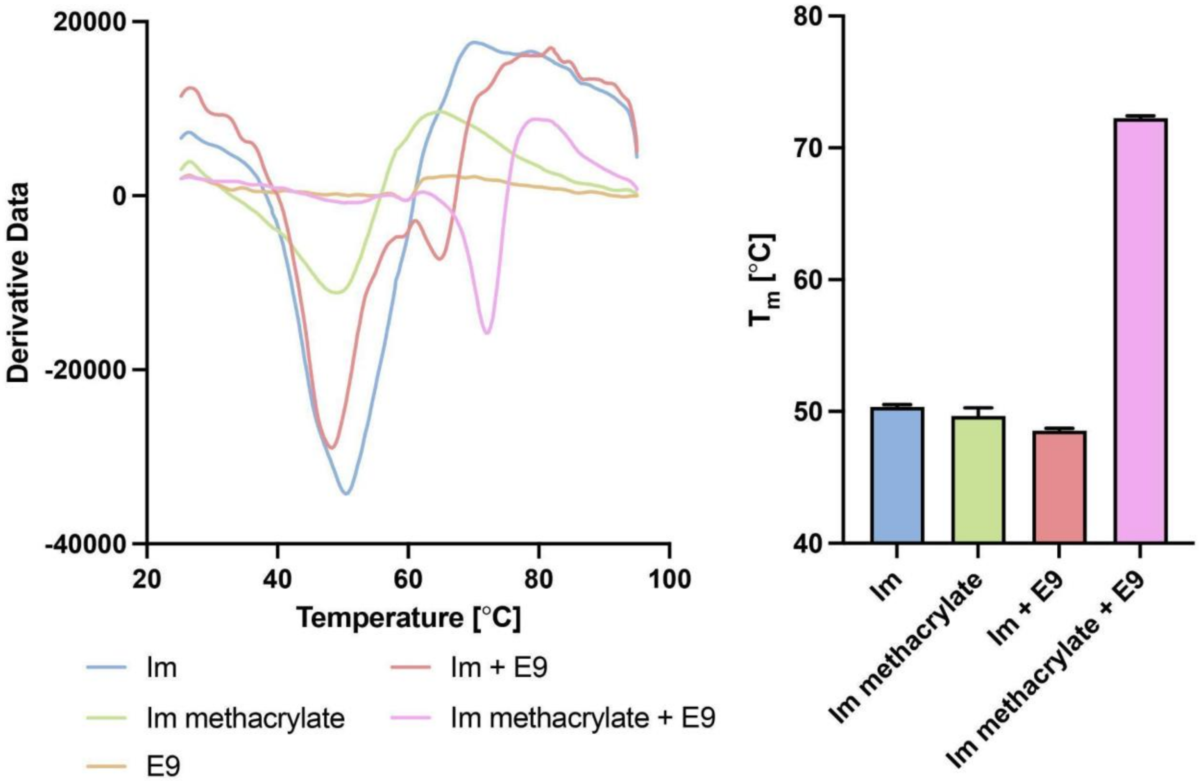

**Supplementary Figure 11. Covalent complex formation significantly stabilizes complex thermal stability.** Left: Average derivative data from 3 samples is shown for each condition. Right: measured melting temperature (calculated from maximum derivative) for different constructs. Error bars represent standard deviation (n=3). Melting temperatures could not be calculated for E9 constructs alone which displayed no clear melting curve.

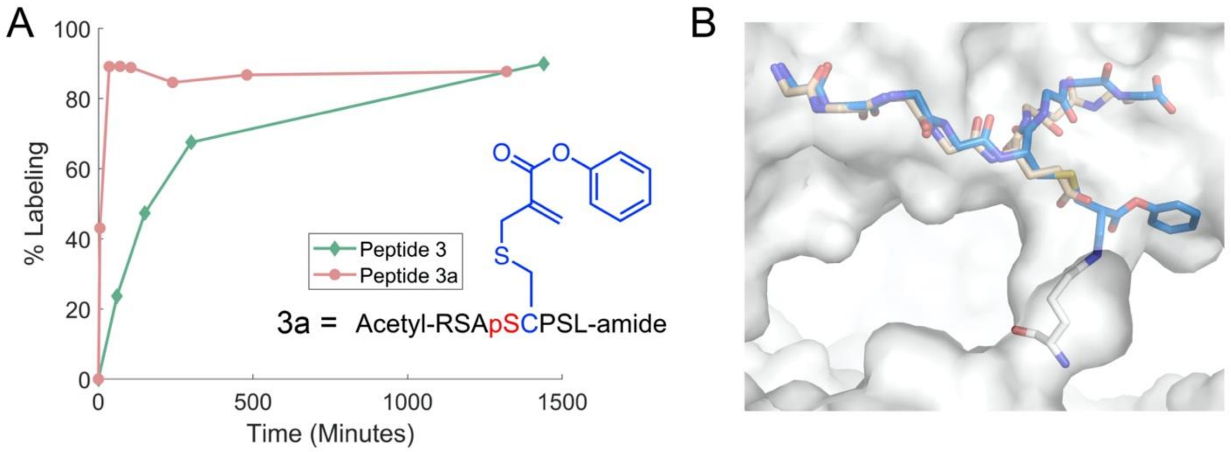

**Supplementary Figure 12. Alternative electrophiles can tune the reaction kinetics. A.** Labeling kinetics of 14-3-3σ (2 µM) by peptide **3** and phenyl ester derivative peptide **3a** (25°C, 5 µM peptide). **B.** CovPepDock model of peptide **3a** bound to 14-3-3σ (blue) overlaid on the structure of parent noncovalent peptide (teal).

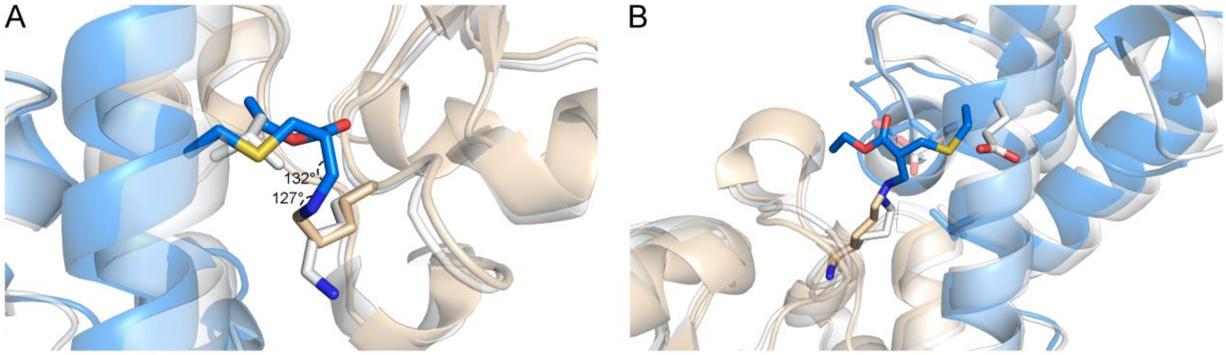

**Supplementary Figure 13. Structural insights provided by CovPepDock for covalent protein reagents. A.** Top-scoring model of the E9-Im9 complex when mutating Val34 of Im9 to the methacrylate warhead. While this position is located in close proximity to the target Lys97 (Cα-Cα distance = 7.5Å), none of the 10 top-scoring models had constraint score < 2, indicating a non-ideal covalent bond geometry. **B.** Docking model (RMSD = 0.758Å) of the E9-Im9 complex when mutating Glu42. While this position seems to be pointing away from the target Lys97, CovPepDock revealed a potential shift of the helix that enables the formation of the covalent bond.

#### Supplementary Tables

##### Supplementary Table 1. Modeling statistics for incorporation of an electrophilic residue to 14-3-3 binding peptides.

| PDB | 14-3-3 Target Lysine | Peptide Position | Best interface backbone RMSD (Å)[a] |
|---|---|---|---|
| 3MHR | 49 | 124 | - |
| | | 125 | 2.197 |
| | | 126 | 1.276 |
| | | 127 | 0.574 |
| | | 128 | 0.978 |
| | | 129 | 0.498 |
| | | 130 | 0.608 |
| | | 131 | 0.609 |
| | | 132 | 1.046 |
| | 122 | 126 | 3.279 |
| | | 127 | 0.619 |
| | | 128 | 0.647 |
| | | 129 | 2.363 |
| | | 130 | 0.843 |
| | | 131 | 0.948 |
| | | 132 | 0.444 |
| | | 133 | 2.087 |
| 3IQU | 122 | 257 | - |
| | | 258 | 6.085 |
| | | 259 | 0.969 |
| | | 260 | 0.455 |
| 3P1N | 122 | 372 | 4.810 |
| | | 373 | 1.642 |
| | | 374 | 0.438 |
| 4IEA | 122 | 620 | - |
| | | 621 | 1.279 |
| | | 622 | 0.529 |
| | | 623 | 1.920 |
| | | 624 | 0.747 |
| | | 625 | 1.068 |
| 4QLI | 122 | 175 | - |
| | | 176 | 3.688 |
| | | 177 | 1.428 |
| | | 178 | 0.640 |
| | | 179 | 0.770 |
| | | 180 | 1.411 |
| 7NFW | 122 | 592 | - |
| | | 593 | 3.979 |
| | | 594 | 0.654 |
| | | 595 | 0.317 |

[a] Best RMSD among the top-10 scoring models that also have a constraint score < 2. If no RMSD is listed it means that no models with constraint score < 2 were generated.

**Supplementary Table 2. Data collection and refinement statistics for 14-3-3σ bound to 3 (PDB: 8C2E) and 8 (8C2F)**

| **PDB** | **8C2E** | **8C2F** |
|---|---|---|
| Protein | 14-3-3σDc | 14-3-3σDc |
| Peptide | **3** | **8** |
| Beam | DESY p11 | DESY p11 |
| <u>*Data Collection*</u> | | |
| Wavelength (Å) | 1.03322 | 1.03322 |
| Space Group | C 2 2 21 | C 2 2 21 |
| Cell Dimensions | | |
| *a, b, c (Å)* | 82.5, 111.7, 63.1 | 82.8, 112.2, 63.0 |
| *α, β, γ (°)* | 90, 90, 90 | 90, 90, 90 |
| Resolution (Å) | 55.92 – 2.20 (2.27-2.20) | 56.14-2.30 (2.38-2.30) |
| I / σ(I) | 12.1 (6.6) | 8.2 (5.0) |
| Completeness (%) | 99.7 (99.4) | 99.9 (100) |
| Redundancy | 12.8 (10.4) | 13.1 (13.1) |
| $CC_{1/2}$ | 0.991 (0.895) | 0.980 (0.720) |
| <u>*Refinement*</u> | | |
| Number of Reflections | 15133 | 13362 |
| $R_{work}$ / $R_{free}$ | 0.175/0.221 | 0.201/0.267 |
| Number of Atoms | | |
| *Protein* | 3516 | 3515 |
| *Ligand/Ion* | 105/4 | 150/4 |
| *Water* | 110 | 59 |
| B-factors | | |
| *Protein* | 18.98 | 17.45 |
| *Ligand/ion* | 18.01/43.35 | 14.18/36.09 |
| *Water* | 23.72 | 18.83 |
| R.M.S deviations | | |
| *Bond Lengths (Å)* | 0.0091 | 0.0072 |
| *Bond Angles (°)* | 1.515 | 1.397 |
| Ramachandran | | |
| *Favored (%)* | 98.69 | 97.39 |
| *Outliers (%)* | 0.00 | 0.87 |
