## Supplementary File 1 - Purified peptides data for "A simple method for developing lysine targeted covalent protein reagents"

Peptide 1:

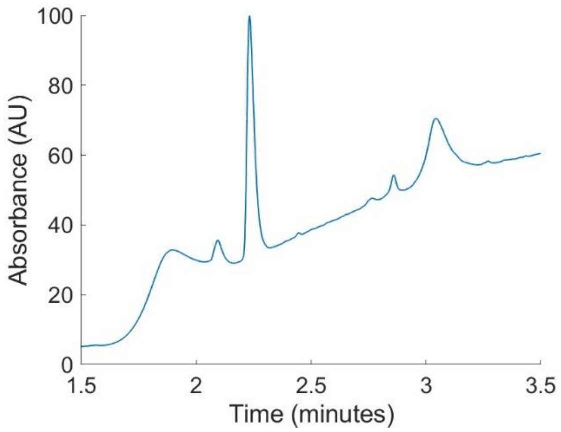


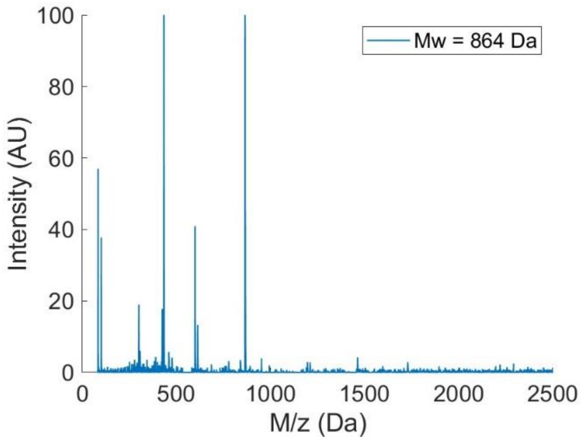


Peptide 2:

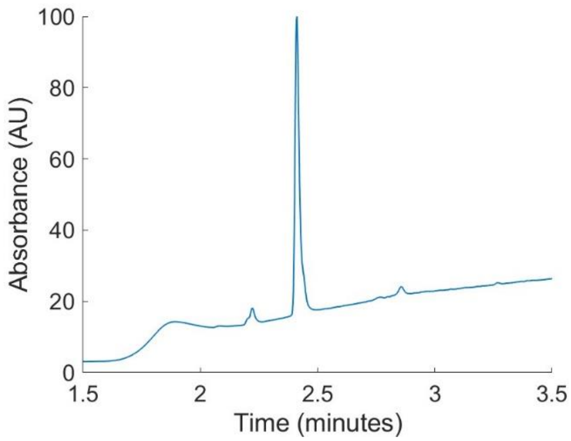


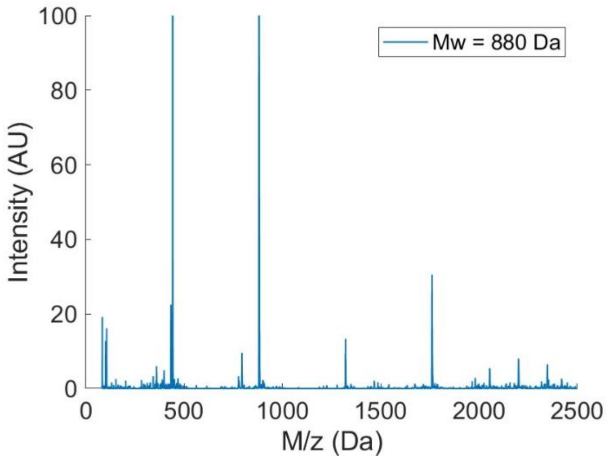


Peptide 3:

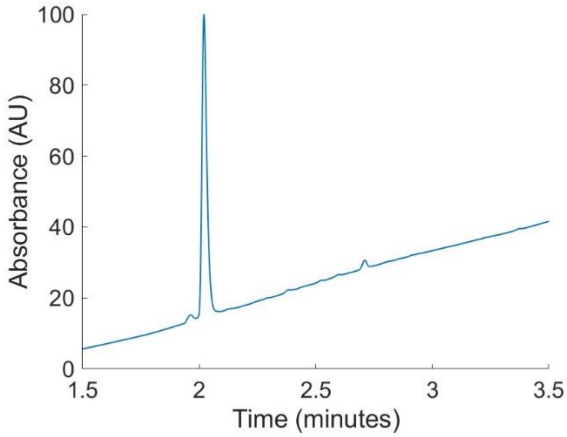


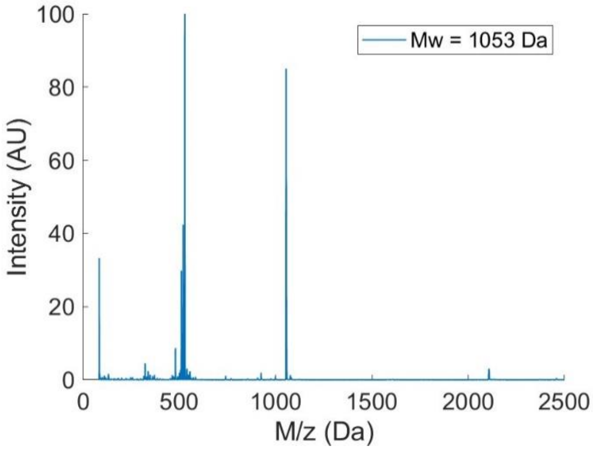

Peptide 4:

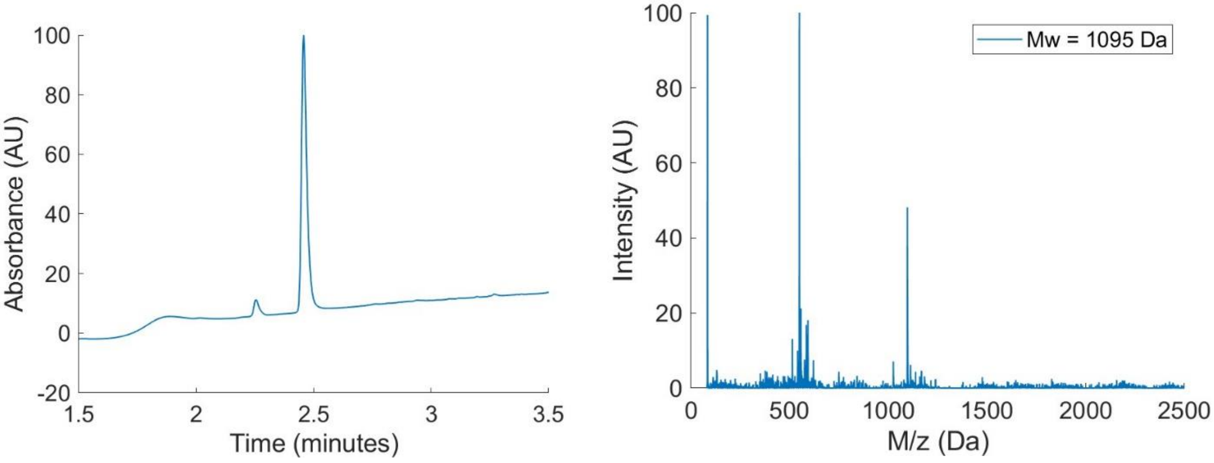


Peptide 5:

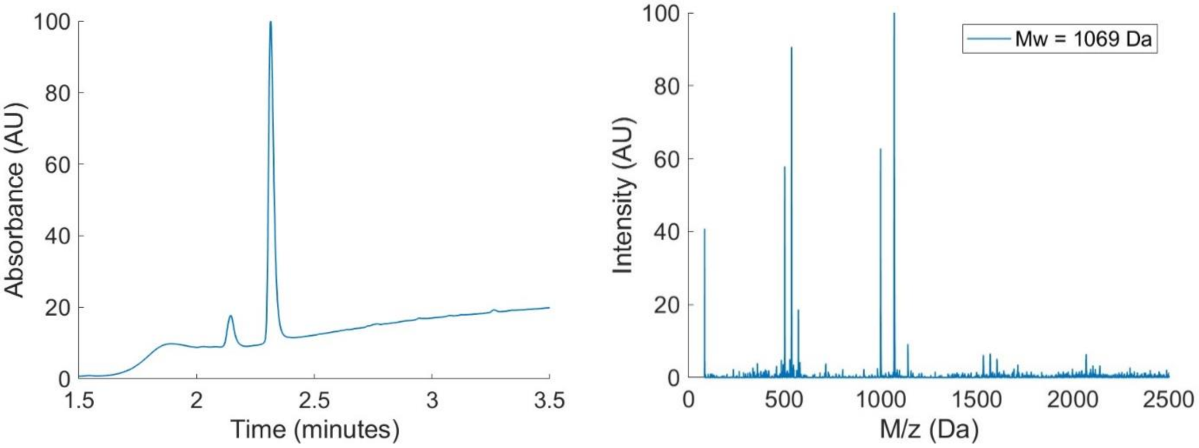


Peptide 6:

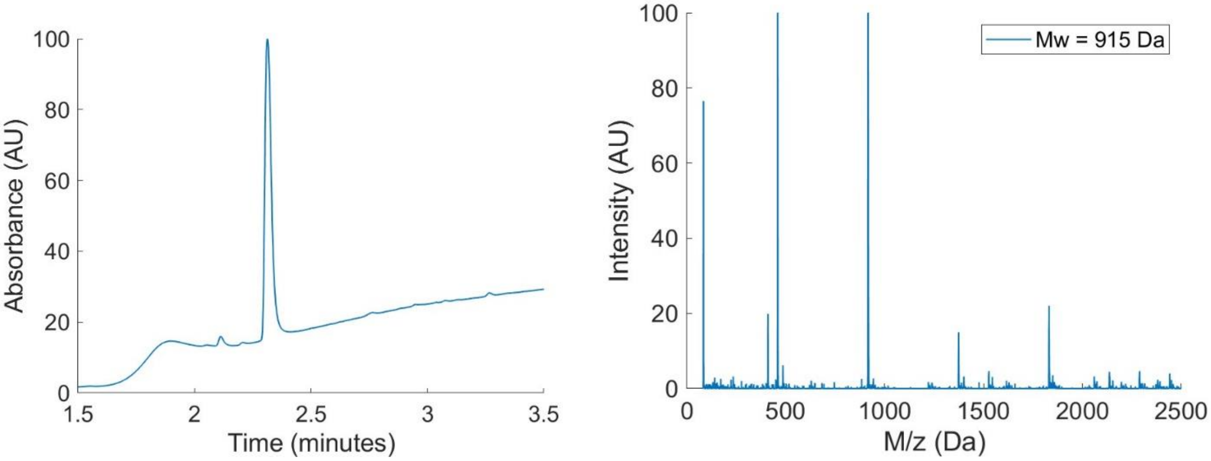

Peptide 7:

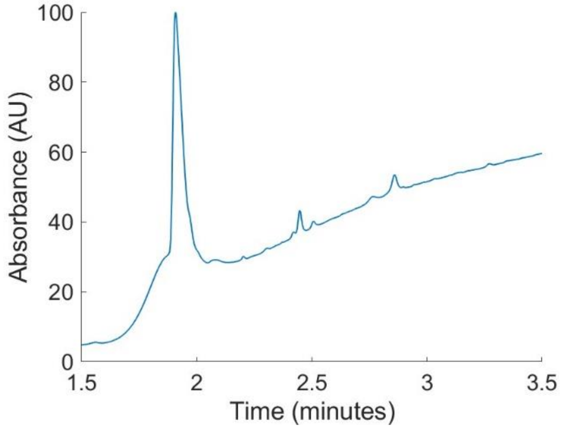


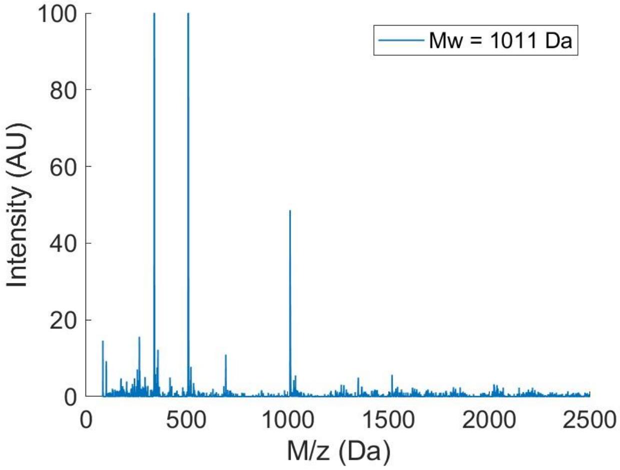


Peptide 8:

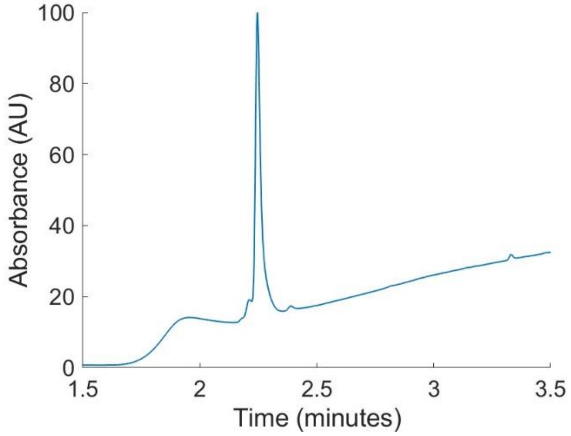


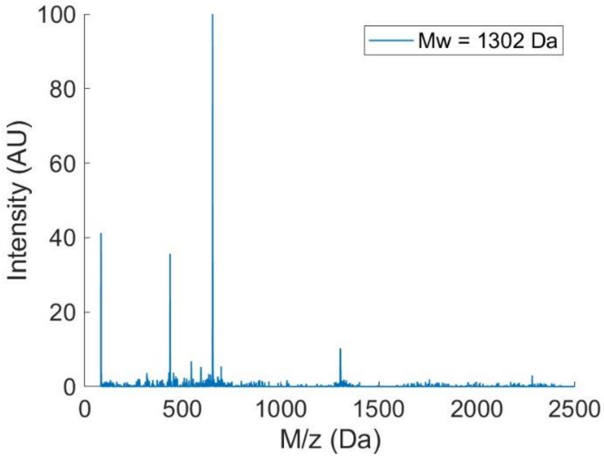


Peptide 9:

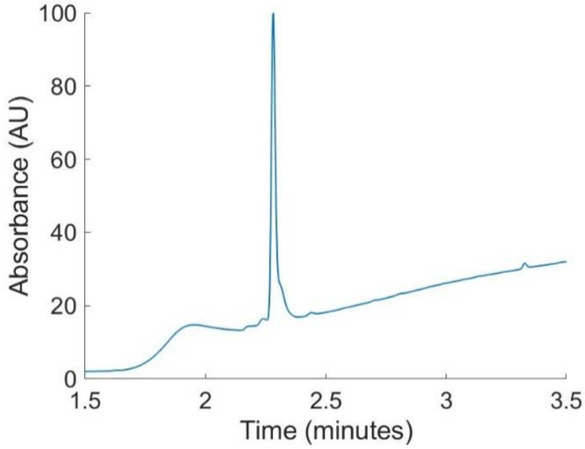


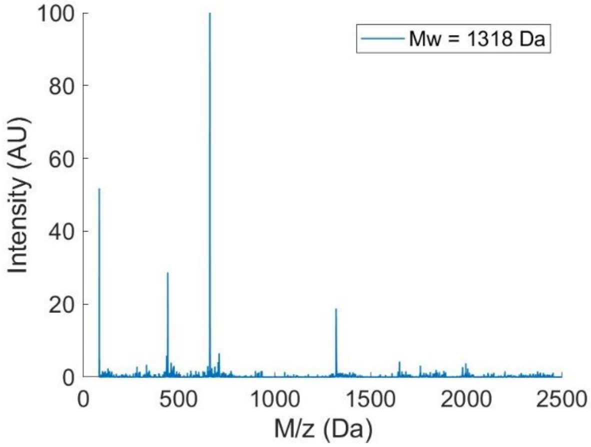

Peptide 10:

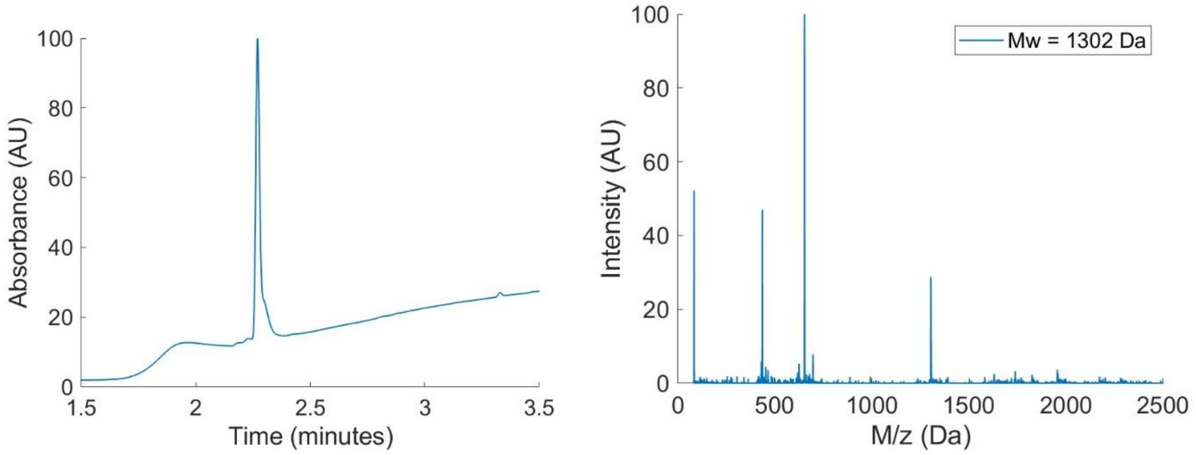


Peptide 11:

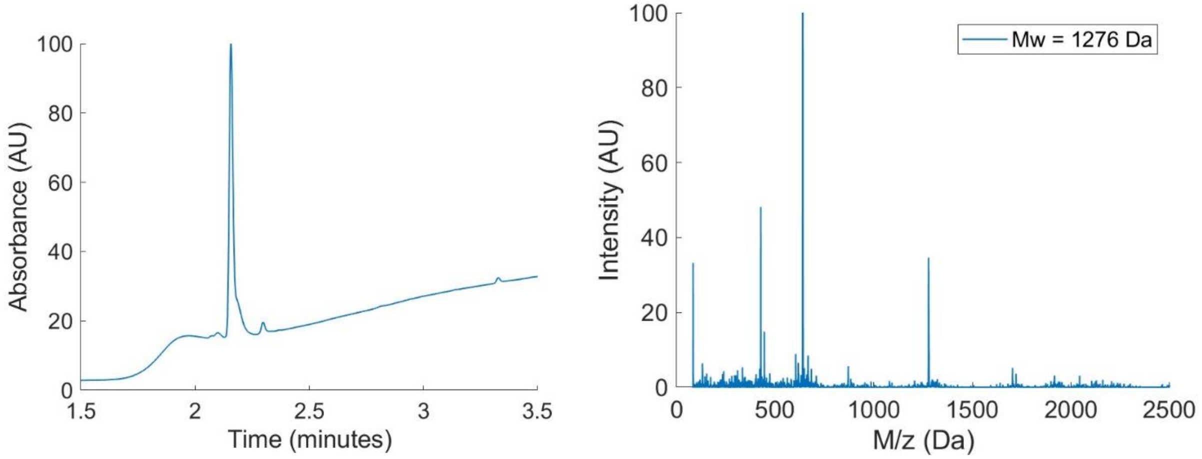


PropargylGly-peptide 3:

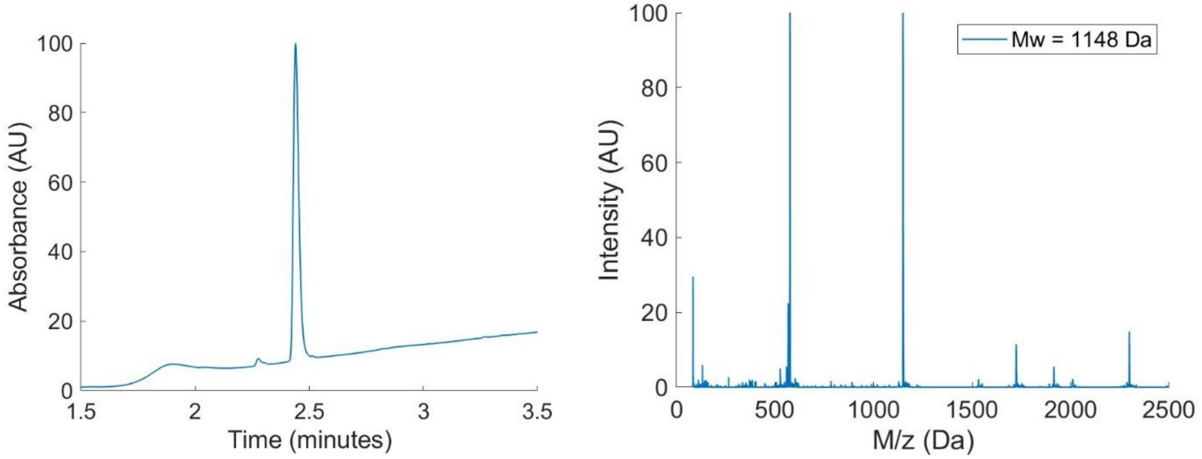

PropargylGly-peptide 8:

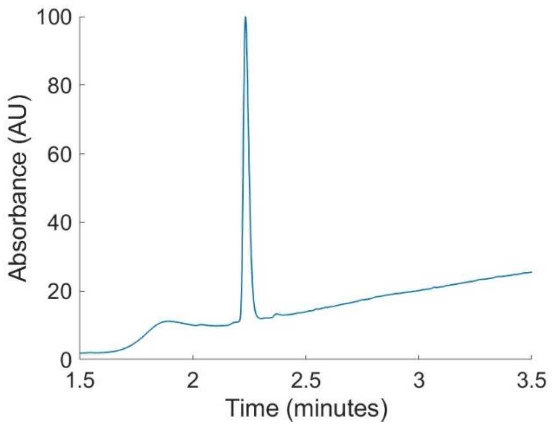


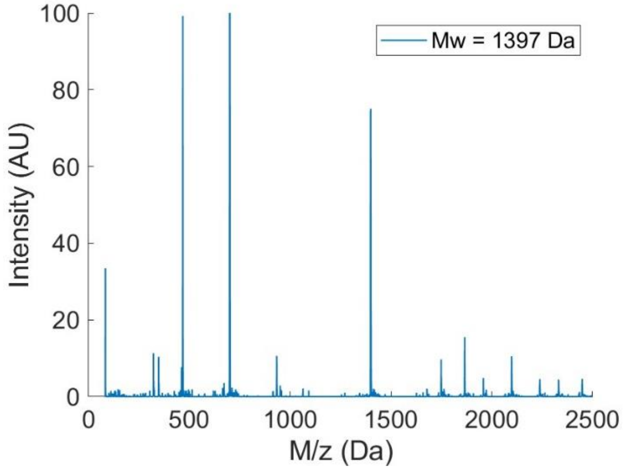


BDP-peptide 3:

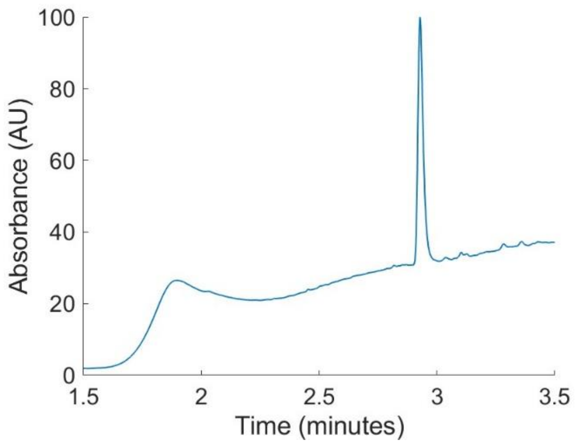


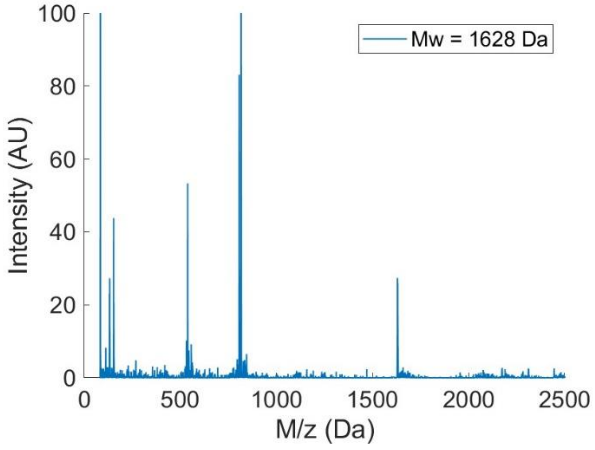


BDP-peptide 8:

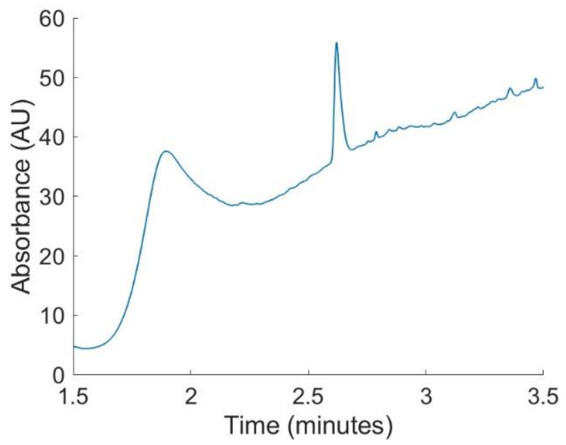


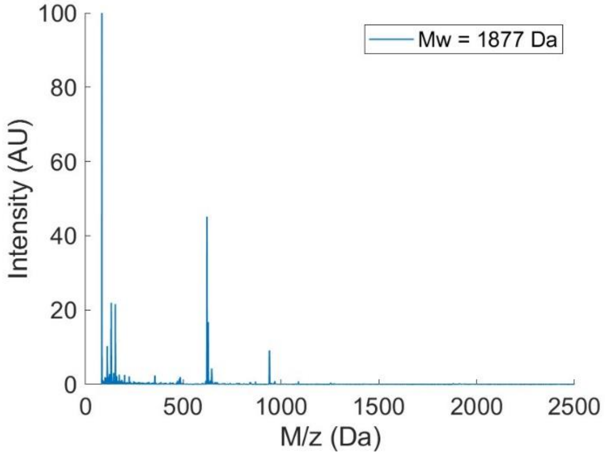

Biotin-Peptide 3:

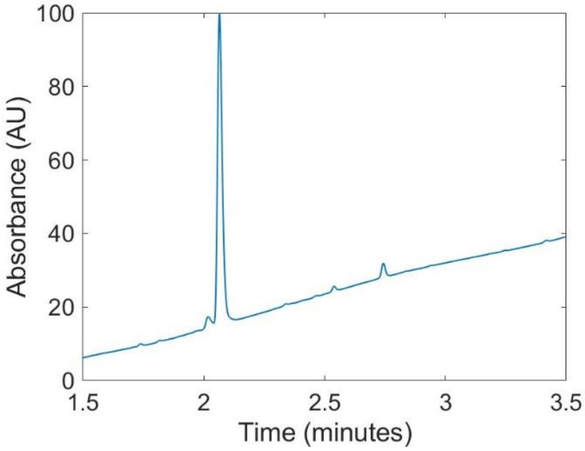


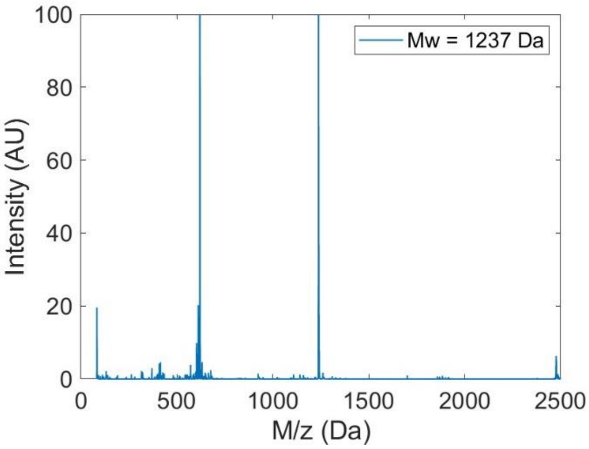


Biotin-Peptide 8:

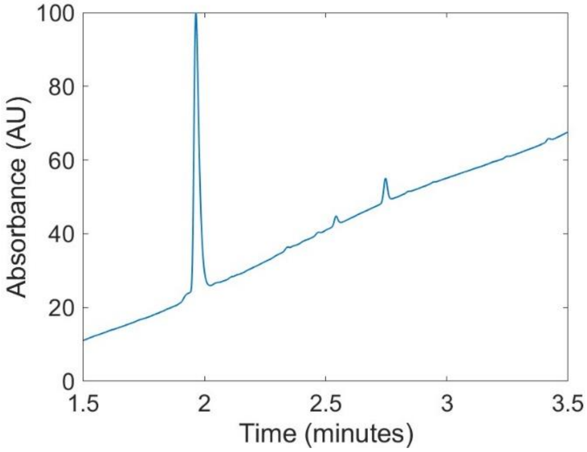


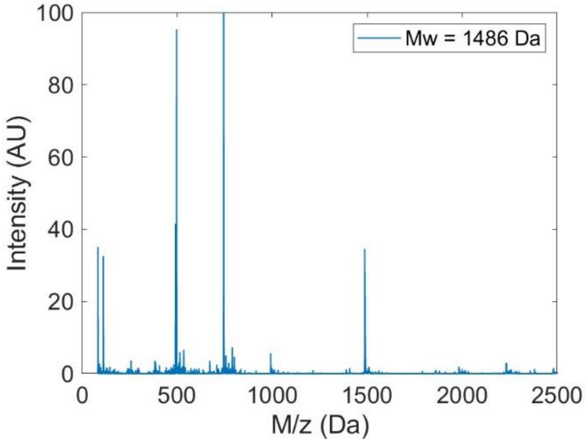
